# ShenQi Compound improves diabetes by modulating pancreatic mitochondrial energy metabolism in GK rats

**DOI:** 10.1101/2025.03.01.640974

**Authors:** Jiushu yuan, huixuan zhang, yulin leng, mingyuan fan, Hanyu liu, hong gao, hongyan xie, haipo yuan, chunguang xie

## Abstract

**Purpose:** ShenQi Compound (SQC) is a traditional herbal formula that has long been utilized in the treatment of type 2 diabetes and its complications. The purpose of this study was to investigate the effect of SQC on mitochondrial energy metabolism in pancreatic tissues of spontaneously type 2 diabetic Goto-Kakizaki (GK) rats.

**Methods:** GK rats were induced into a diabetic model using a high-fat diet. They were randomly divided into 3 groups (n=8): diabetes model group, SQC group (14.4 g/kg/d) and metformin (Met) group (0.1 g/kg/d). Another 8 Wistar rats were taken as controls. Weight, Blood glucose was monitored weekly in each group,After 12 weeks of gavage. Fasting blood glucose and lipid levels were evaluated, and histopathological changes in the pancreas were assessed by hematoxylin-eosin (HE) staining, serum fasting insulin(FINS) and pancreatic mitochondrial respiratory chain complex I-V (complexI-V) levels were measured by enzyme-linked immunosorbent assay (ELISA), reactive oxygen species ((ROS)) levels were detected by chemiluminescence, superoxide dismutase (SOD) levels were detected by xanthine oxidase, and glutathione (GSH) and adenosine triphosphate (ATP) levels were detected by colorimetric assay. And the uncoupling protein(UCP-2) protein and mRNA expression levels were detected by protein blotting and real-time quantitative PCR.

**Results:** SQC treatment significantly decreased (P<0.01) the levels of FBS, FINS,triglyceride s(TG), total cholesterol (TC), and low-density lipoprotein cholesterol (LDL-C) (P<0.01) and significantly increased (P<0.01) the levels of denser lipoprotein cholesterol (HDL-C) in GK rats. Pancreatic histopathological damage was improved after SQC treatment, and complexI to complexIV significantly decreased (P<0.05) and complexV, SOD and GSH significantly increased (P<0.05) in pancreatic tissues. In addition, SQC reduced the content of ROS and ATP in pancreatic tissues (P<0.01) and down-regulated UCP-2 protein and mRNA expression (P<0.01).

**Conclusion:** SQC improves glucose-lipid metabolism and attenuates pathological damage of pancreatic tissues in T2DM rats, possibly by regulating mitochondrial energy metabolism in pancreatic tissues. The effects of SQC are not well understood. These findings provide new insights into the mechanism of action of SQC in the treatment of T2DM and its associated neurodegenerative effects.

## Introduction

The prevalence of diabetes mellitus has been steadily increasing in recent years, and diabetes mellitus, as one of the major public health problems in the world in the 21st century, is expected to reach 643 million by 2030 and 783 million by 2045 [1,2]. As diabetes progresses, cardiovascular and cerebrovascular diseases, blindness, renal failure, gangrene and even death may occur [3], bringing serious threats to patients’ physical and mental health while consuming a large amount of medical and healthcare resources, and imposing a heavy burden on society [4,5]. Currently, the treatment of T2DM is mainly based on oral hypoglycemic agents and improvement of insulin resistance, however, there are problems such as adverse reactions and poor efficacy [6], therefore, there is an urgent need to find effective targets and drugs to solve this problem.

Recent studies have shown [7] that the emergence of T2DM is closely related to abnormalities in pancreatic mitochondrial energy metabolism. Under physiological conditions, pancreatic islet β-cells couple mitochondrial oxidative phosphorylation and ATP production to achieve normal insulin secretion, and the reduced nicotinamide purine (NADH) and succinic acid (FADH2) produced during this process gradually transfer electrons through the mitochondrial respiratory chain accompanied by the gradual release of energy, i.e., ATP [8,9]. ATP not only provides energy for cellular metabolism, but also serves as a signal for regulation of insulin secretion and fine regulation of insulin secretion [10]. In addition to providing energy for cellular metabolism, ATP also serves as a signal for controlling and fine-tuning insulin secretion[10,11]. However, prolonged elevation of blood glucose leads to over-operation of the mitochondrial respiratory chain, causing electron leakage from the respiratory chain, and the leaked electrons are directly handed over to O2 at the end of the respiratory chain, thus generating excessive ROS, which aggravate cellular damage and even mediate cell apoptosis [12–17]. On the other hand, it can also up-regulate UCP-2 on the inner membrane of mitochondria, affecting the normal secretion of insulin [18]. It is clear that safeguarding pancreatic β-cell mitochondrial function is a crucial step toward improving pancreatic-cell functionality, maintaining blood glucose stability, and serving as a vital link in the chain of events that effectively postpones the advancement of diabetes mellitus.

SQC is a traditional Chinese medicine compound formula widely used in the treatment of T2DM and its complications, which consists of 8 Chinese medicines: *Panax ginseng C.A.Mey, Astragalus mongholicus Bunge, Rehmannia glutinosa, Dioscorea oppositifolia L, Trichosanthes baviensis Gagnep, Cornus officinalis Siebold & Zucc, Salvia miltiorrhiza Bunge, Rheum palmatum L*.and a large number of previous clinical studies have confirmed that SQC has the effects of improving insulin resistance, regulating glucose and lipid metabolism,and improving the function of pancreatic islets [19–21]. Its benefits are significantly clearer in islet cell protection, and previous animal investigations have confirmed this. SQC was also shown in previous animal studies to protect pancreatic islet cells by enhancing the β-cell resistance to damage in diabetic blood glucose fluctuation model rats [22] and inhibiting apoptosis of pancreatic islet β-cells in GK rats [23,24]. However, its specific role has not been fully elucidated, so we hypothesize that the protection of pancreatic islet cells by SQC in diabetic rats may be achieved by improving the metabolism of mitochondrial metabolic capacity of pancreatic tissues. Based on this, the present study was to further investigate the protective effect of SQC on mitochondrial energy metabolism function in pancreatic tissues of diabetic GK rats on the basis of the previous study, and to provide new therapeutic evidence for the clinical use of SQC in the treatment of T2DM.

## Materials and Methods

### Drug Preparation

Chinese medicine ginseng and astragalus compound formula consists of ginseng 15g; astragalus 15g; Radix et rhizoma 10g; pollen of smallpox 10g; Chinese yam 10g; Cornus officinalis 10g; Salviae Miltiorrhizae 10g; systematic rhubarb 6g, Chinese herbal medicine was purchased from Chengdu New Lotus Herb Co. The second and the third time were given 8 times the volume of water, boiled over high heat and then poured the medicinal liquid for 20 minutes, the three poured medicinal liquid was mixed and filtered, and the medicinal liquid was concentrated in a rotary distiller at a temperature of 65 ℃, and concentrated until the concentration of the extract containing the raw medicine was 14.4g/ml, poured and took the extract, and preserved it at 4 ℃[25].

Positive control drug is metformin hydrochloride tablets, specification 50mg/tablet×20 tablets/box, Shanghai Squibb Co.

### Animal feeding, grouping and treatment

This study was approved by the Experimental Animal Ethics Committee of the Affiliated Hospital of Chengdu University of Traditional Chinese Medicine. Twenty-four 7 - 9-week-old male GK rats weighing 220-280 g and eight 7 - 9-week-old male Wistar rats weighing 250-270 g were purchased from Shanghai Slaughter Laboratory Animals Limited Liability Company (Laboratory Animal License No.: SCXK(Shanghai) 2017-0005). The animals were housed in an SPF-rated laboratory of the Sichuan Academy of Traditional Chinese Medicine (SACTM) at a temperature of 22°C, relative humidity of 50%, air circulation, and 12 hours of alternating illumination.

After all rats were acclimatized according to 4 rats/cage for 7 days, the GK rats were randomly divided into the model group (DM), metformin group (Met), and sennaqi compound group (SQC) according to the principle of randomization with 8 rats in each group, and 8 Wistar rats were set up as a blank control group (NC), and the diabetic models of the GK rats were prepared by giving the high-fat chow to the GK rats continuously fed (regular chow 88.2%, refined lard 10%, cholesterol 1.5%, and porcine bile salt 0.3%), while the Wistar rats were given regular chow (71% carbohydrate, 4.5% fat, and 20% protein).

Based on random blood glucose values >11.1 mmol/L, the rats were successfully modeled, and the drug administration began. The normal group and the model group were gavaged with saline at a dose of 5 mLkg-1 d-1; the ginseng qiqi formula group was gavaged with ginseng qiqi formula suspension at a dose of 14.4 g·kg-1·d-1 (equivalent to 10 times of the dose of the adult, and for the adult based on a body mass of 60 kg, each 1 mL of the infusion contained 1.44 g of raw herbs); the metformin group was gavaged at 0.1 g/(kg-d); the body mass of rats was measured regularly every week, and the amount was adjusted according to changes in body mass, and the drug was administered for 12 consecutive weeks.

### Measurement of body weight fasting blood glucose (FBG) levels

Every Monday, the rats were fasted from 9:00 a.m. and weighed one by one after 5:00 p.m. (8 h), then blood was collected from the tail tip and the FBG level was measured with a glucose meter.

### Lipid Profile Test

After 12 weeks of drug intervention,Blood was collected from the abdominal aorta of rats after a 10-h fast, and TC,TG, LDL-C, and HDL-C biochemistry kits were purchased from Nanjing Jianjian Biological Company Limited, with the following article numbers: A111-1-1, A110-1-1, A113-1-1, and A112-2, respectively, and the serum concentrations of TC, TG, LDL-C, and HDL-C were measured using a full-automated biochemistry analyzer (Radiometer, Shenzhen, China) according to the manufacturer’s protocol.

### Elisa assay for serum insulin and pancreatic tissue mitochondrial respiratory chain complex I-V (complexI-V) levels

Pancreatic tissues were rapidly isolated after blood sampling and centrifugation. The pancreatic tissues were analyzed by the corresponding ELISA kits [rat FINS (Shanghai Zuo Cai Biotechnology Co., Ltd., ZC-54525), rat complexI (MM-70671R2), complexII (MM-70264R2), complexIII (MM-71168R2), complex IV (MM-7066R2), complex *Ⅴ* (MM-71172R2), mitochondrial respiratory chain complex kit were purchased from Wuhan Saipai Biotechnology Co.), complex V (MM-71172R2), mitochondrial respiratory chain complex kit were purchased from Wuhan Saipei Biological Co., Ltd.] to detect the levels of FINS, complexI, complexII, complexIII, complexIV and complexV, respectively, and all the procedures were strictly in accordance with the instructions of the kits.

### HE staining for detection of pancreatic histopathology

Pancreatic tissues were fixed in 4% paraformaldehyde at room temperature for 24 h. After the steps of dehydration, trimming, embedding, sectioning (4 μm thick), hematoxylin and eosin staining, and sealing, the slices were subjected to image acquisition using a Pannoramic 250 digital slice scanner (3DHISTECH (Hungary)).

### Detection of oxidative stress indicators and ATP content in pancreatic tissues

The ROS, SOD, GSH and ATP contents in pancreatic tissues were determined using ROS fluorescence test kit (E-BC-K138-F), SOD typing test kit (E-BC-K022-M), GSH colorimetric kit (E-BC-K030-M), and ATP colorimetric kit (E-BC-K157-M), respectively, which were purchased from Wuhan Elite Bio-Technology Co.

### Quantitative Real-Time PCR Assays

The total RNA of pancreatic tissue was extracted using TRIzol reagent (Foregene), and the RNA concentration and purity were detected by K2800 Nucleic Acid Analyzer (Beijing Kai-O Technology Development Co., Ltd.), Real-time PCR kit (ABM Canada) was carried out in accordance with the instructions, and the PCR system was used for the amplification (the PCR reaction conditions were 95 ℃ pre-denaturation for 100 s, 95 ℃ denaturation for 15 s, 60 ℃ annealing for 55 s, 42 cycles), and the primers were quantified afterward. The primers were synthesized by Sangong Bioengineering (Shanghai) Co., Ltd. and the primer sequences were UCP-2 upstream 5’-CAGATGTGGGTAAAGGTCCGCTTCC-3’, downstream 5’- TCGTGCAATGGTCTTGTAGGCTTC-3’, GADPH upstream 5’- TCGTGCAATGGTCTTGTAGGCTTC-3’, and GADPH upstream 3’, and GADPH upstream 3’. 3’, GADPH upstream 5’-CAGATGTGGTAAAGGTCCGCTTCC-3’ downstream, 5’- TCGTGCAATGGTCTTGTAGGCTTC-3’ using 2-ΔΔΔTCTTAGGCTTC-3’. The relative expression of target genes was calculated by the 2-ΔΔct method [26].

### Western Blotting Assays

A homogenate of pancreatic tissue was made by taking 0.1 g of fresh pancreatic tissue and placing it in pre-cooled PBS, adding 450 κL of saline, and grinding it thoroughly on ice.Total proteins were extracted from pancreatic tissues using RIPA lysate (biosharp), proteins were quantified using BCA kit (biosharp), and equal amounts of proteins were separated by SDS-PAGE and transferred to polyvinylidene difluoride membranes. The membrane was closed with skimmed milk for 1 hour. The membrane was incubated with primary antibody UCP-2 (1:1500, Affinity, USA, AF7018), GAPDH (1:10 000 Affinity, USA, AF7021) at 4 ℃ overnight; the membrane was washed with TBST buffer and then the membrane was incubated with secondary antibody (1:10000, Hangzhou Lianke Bio-Technology Co. Ltd, 70-GAM0072) for 1 h at room temperature. After the membrane was washed with TBST buffer and secondary antibody (1:10000, Hangzhou Lianke Biotechnology Co., Ltd., 70-GAM0072) was added, the membrane was exposed to the luminescent solution mixed with ECL A and ECL B in equal proportions. GAPDH was used as an internal reference, and the relative protein expression was analyzed in gray scale by using ˂Qinxiang Chemical Analysis Software˃, and the relative protein expression was determined by the gray scale value of the target protein/internal reference protein bands.

## Statistical methods

Data were expressed as mean ± standard deviation (SD) and analyzed by one-way ANOVA followed by the LSD t-test. All statistical analyses were performed using GraphPad Prism Software (version 9.5.0), with P values < 0.05 considered statistically significant.

## Results

### Effect of ginseng and SQC formula on body weight, FBG and FINS in rats

As mentioned above, blood glucose ≥11.1 mmol/L was regarded as successful modeling of diabetes, and then the SQC and Met groups were fed 14.4 g/kg/d for SQC and 0.1 g/kg/d for Met according to the doses of previous studies [27–29], and the Normal and Diabetes groups were fed with saline for 12 weeks, and the FBG of rats in each group was monitored during the experiment.

From the results of Fig. 1A, it was found that the body weight level of the model group decreased at 0, 4, 8, and 12 weeks compared with the normal group (P ˃ 0.05), and after drug treatment, the body weight level of the SQC group and the Met group decreased compared with the model group, but none of them was statistically significant, and from Fig. 1B, it was found that the body weight of the various rats showed a tendency to increase as the experiment progressed (P ˃ 0.05), At 4 weeks, the body weights of rats in each group were close to the same, and at 8 and 12 weeks, the body weights of rats in the SQC group were lower than those in other groups (P ˃ 0.05), which showed that the effect of SQC on the body weights of GK rats was not obvious.

**Figure 1.**
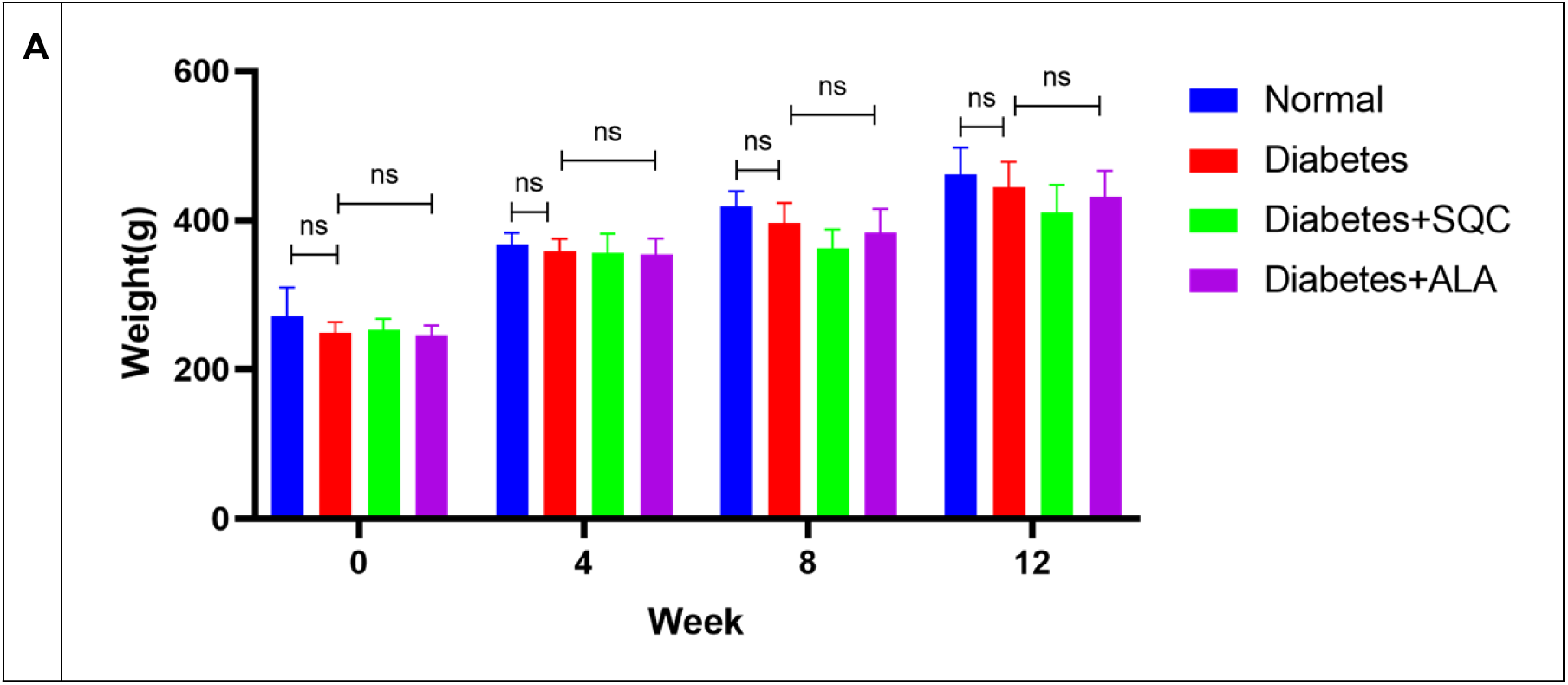
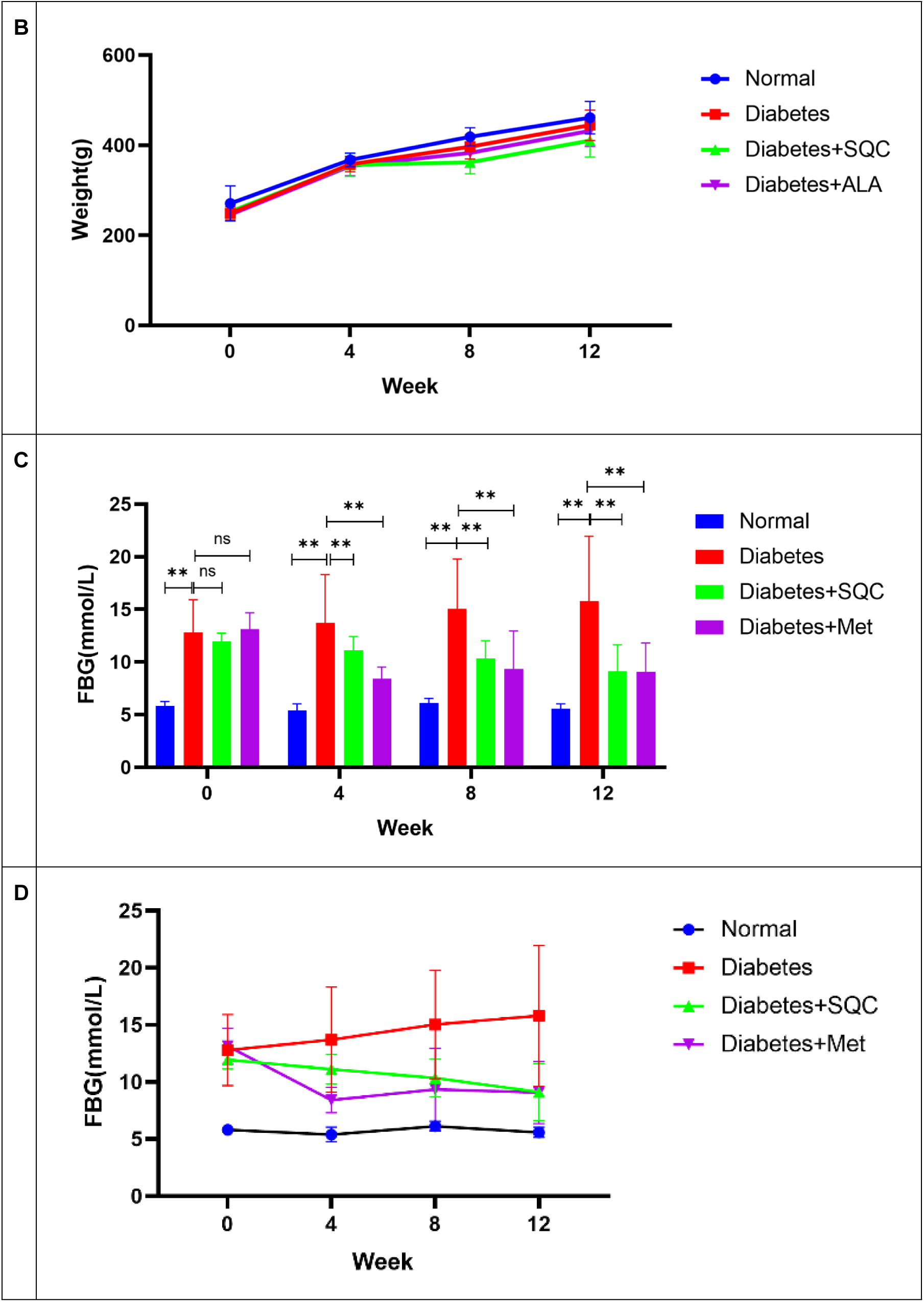
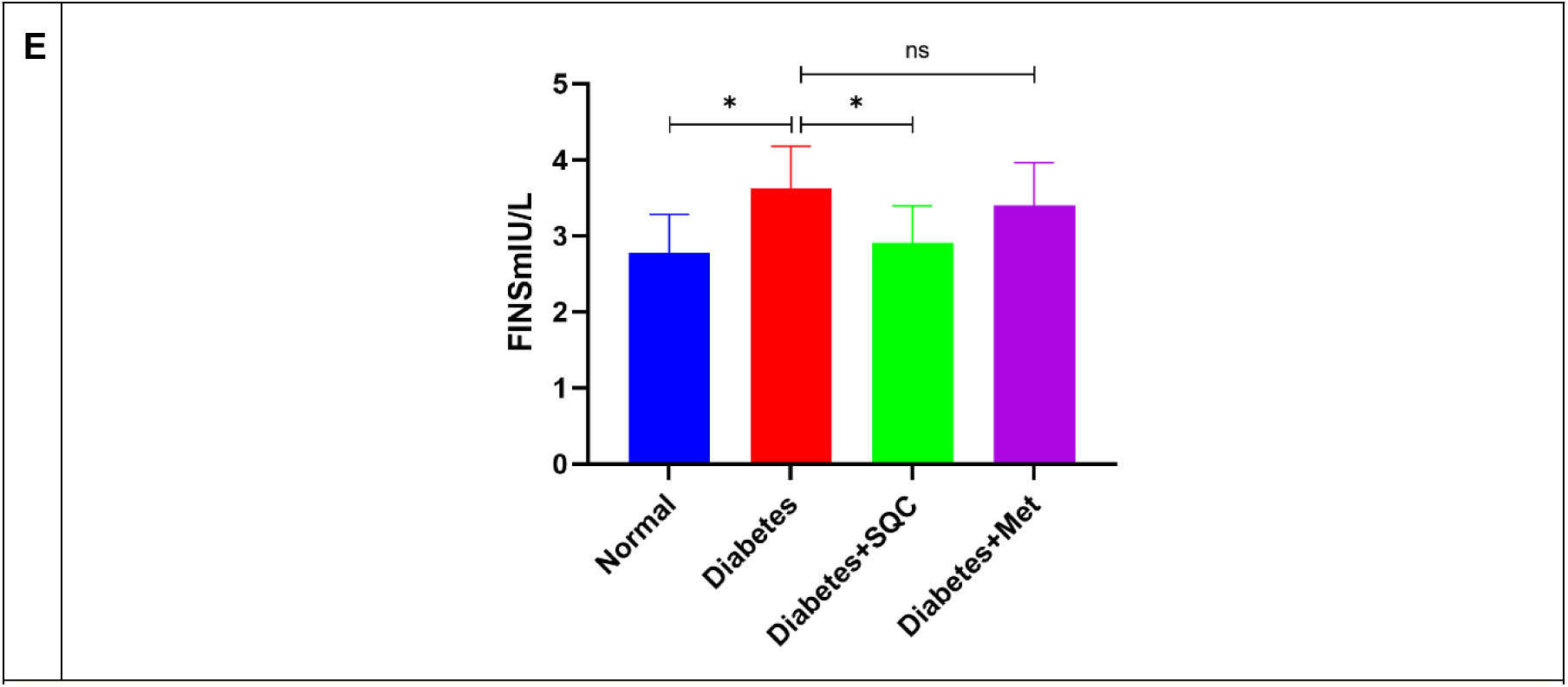
Effects of SQC and Met on Wight*、*FBG and FINS in GK rats. (A, B)Effects of SQC and Met on Wight in GK rats,(C, D) Effects of SQC and Met on FBG in GK rats, (E) Effects of SQC and Met on FINS in GK rats. All data are expressed as mean ± SD (n = 6-8). * p < 0.05,** p < 0.01. Abbreviation: NS, not significant.

From the results of Figure 1C, it was found that before drug intervention, the FBG level of GK rats was significantly higher than that of Wiater rats (P<0.01), which was consistent with the characteristics of GK rats, while there was no significant difference in FBG among the Diabetes group, the Met group of rats, and the SQC group; at 4, 8, and 12 weeks of drug intervention, the FBG in the Diabetes group was still significantly elevated when compared with the Normal (P <0.01), and compared with Diabetes group, FBG levels in both SQC and Met groups decreased significantly (P<0.01), which indicated that the hypoglycemic effect of SQC was similar to that of Met.

As can be seen from Figure 1D, as the experiment proceeded, there was no significant change in the FBG of the rats in the Normal group, while the FBG of the model Diabetes group was always at a high level, and with the prolongation of the experiment the FBG gradually increased, and the FBG of the Met group and the SQC group both showed a clear downward trend. 0-4 weeks, the Met showed a significant downward trend compared with that of the SQC, and from 4 to 8 weeks the FBG of the two groups continued to decrease, while the Met group gradually increased and had a fluctuating rebound trend. In the Met group, FBG gradually increased, with a fluctuating rebound trend. At 12 weeks, FBG of rats in the Met and SQC groups was still at a higher level than that of the Normal group, but compared with the model group, FBG was still on a downward trend, and the SQC group was on a par with the Met group. It can be seen that the hypoglycemic effect of SQC was smoother in the Met.

In addition to this, FINS levels were also examined as shown (Fig. 1E), which were significantly higher in the model group compared to the normal group (P < 0.05), and after drug treatment, FINS levels were significantly lower in the SQC group compared to the model group compared to the Met group (P < 0.05)

### Effect of ginseng and SQC formula on blood lipids in rats

To determine whether SQC improves lipids in T2DM rats, we performed biochemical analysis of TC, TG, LDL-C, and HDL-C. Our results showed that the levels of TC, TG, and LDL in the modeling group were significantly higher than those in the normal group (P<0.01), and the levels of HDL-C were significantly lower than those in the normal group (P<0.01); however, after pharmacological interventions, a significant decrease in TC, TG, and LDL was seen in both SQC and Met rats compared to the modeling group (P<0.01), and a significant increase in HDL-C was seen (Figure 2A- D, P<0.01). Taken together, these results suggest that SQC can improve lipid metabolism disorders in diabetic rats.

**Figure 2.**
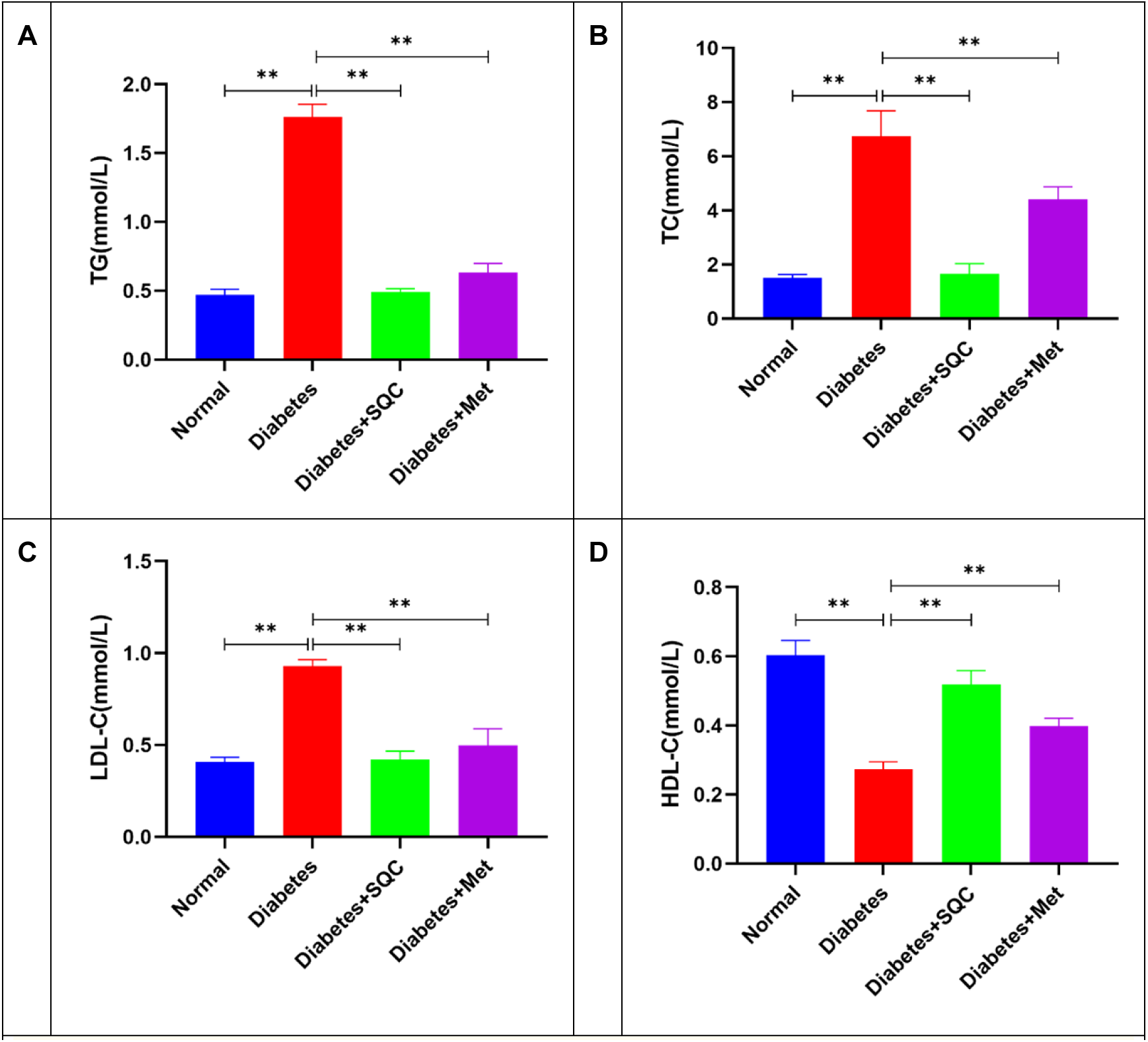
Effects of SQC and Met on lipids in GK rats. (A) Effects of SQC and Met on TG in GK rats, (B) Effects of SQC and Met on TC in GK rats, (C) Effects of SQC and Met on LDL-C, in GK rats, and (D) Effects of SQC and Met on HDL-C in GK rats, all data are expressed as mean ± SD (n = 6). * p < 0.05, ** p < 0.01.Abbreviation: NS, not significant.

### The effect of ginseng and SQC formula on the histopathology of rat pancreas

HE staining results showed that the pancreatic tissue of rats in the Normal group was structurally intact, with clear lobes and no thickening, and all types of cells in the pancreatic islets had normal morphology and clear nuclei (Figure 3A). On the contrary, Diabetes pancreatic islets were mildly atrophied, islets were irregularly shaped, exocrine vesicles invaded the islets (indicated by yellow arrows), the number of pancreatic islet cells was reduced, proliferation of fibrous tissues at the edge or center of the islets was seen (indicated by blue arrows), which separated the pancreatic islet cells, and an increased number of fibroblasts with elongated oval or ovoid nuclei was seen, and a slight hyperplastic vasculature was seen around the islets in the ducts and peripheral inflammatory cell infiltration (indicated by green arrows) as in (Figure 3B). After 12 weeks of pharmacologic intervention, the degree of pathological changes was reduced in both SQC(Figure 3C) and Met groups(Figure 3D), with more pronounced improvement in SQC (Figure 3C) than in the control group.

**Figure 3.**
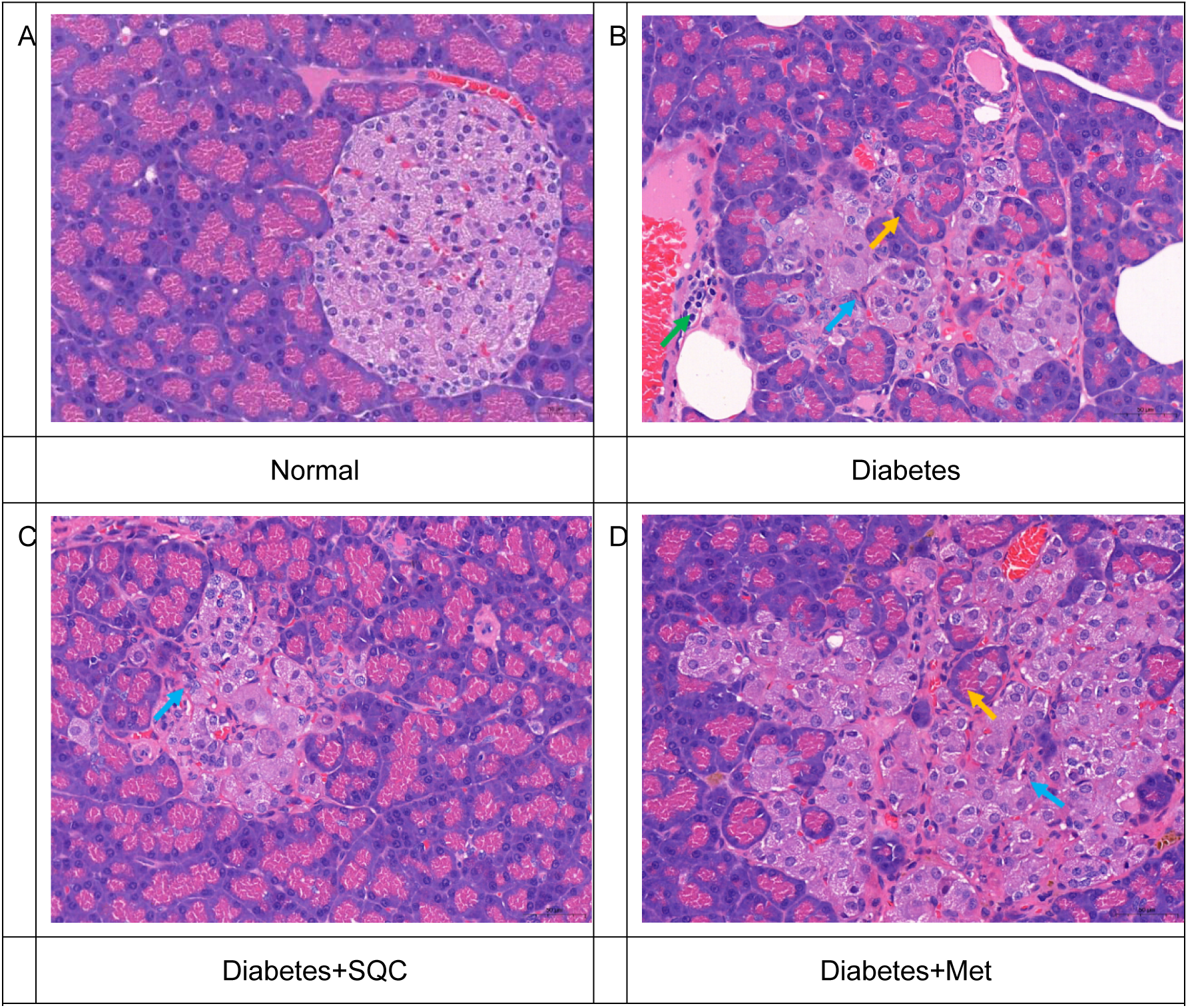
Effect of SQC on histopathology of rat pancreas, HE is staining of pancreas, yellow arrows indicate invasion of adenoids into pancreatic islets, blue arrows indicate proliferation of fibroblasts, and green arrows indicate invasion of lymphocytes, Magnification, 400×; Scale bar=50 μm(3A-D)

### Mitochondrial respiratory chain complexes I-V in pancreatic tissue of SQC rats

Pancreatic β-cell mitochondria are the main site of energy metabolism in pancreatic β-cells, and the effects of SQC and Met groups on the expression of mitochondrial respiratory chain complex I-V in pancreatic tissues of GK rats are shown in Fig. 4A-E. The levels of complex I, complex II, complex III, complex IV, complex V in the model group were significantly higher than that in the control group (P<0.05, P <0.01), however, the levels of complex I, complexⅡ, complexⅢ, complexⅣ, and complexⅤ were significantly down-regulated in the SQC group compared with the model group (P<0.05, P<0.01), and the levels of complexⅠ, complexⅡ, and complexⅣ in the Met group were significantly lower than those in the model group (P< 0.05, P<0.01), and complex III and complex V decreased in the Met group compared with the model group, but without statistical significance. It can be seen that both SQC and Met have the potential to improve the energy metabolism of mitochondrial respiratory chain in diabetic rats, and the efficacy of SQC is better than in Met.

**Figure 4.**
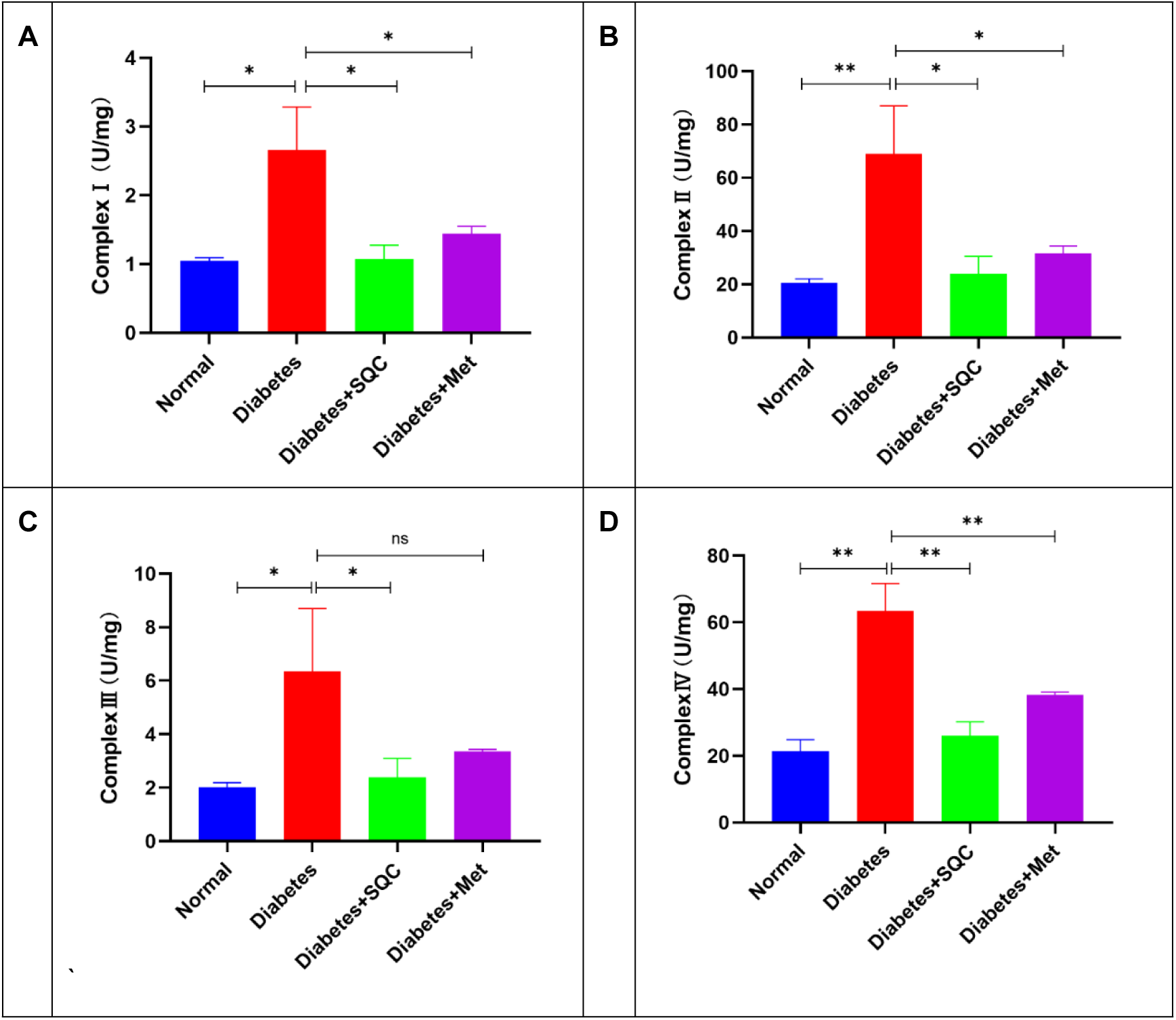
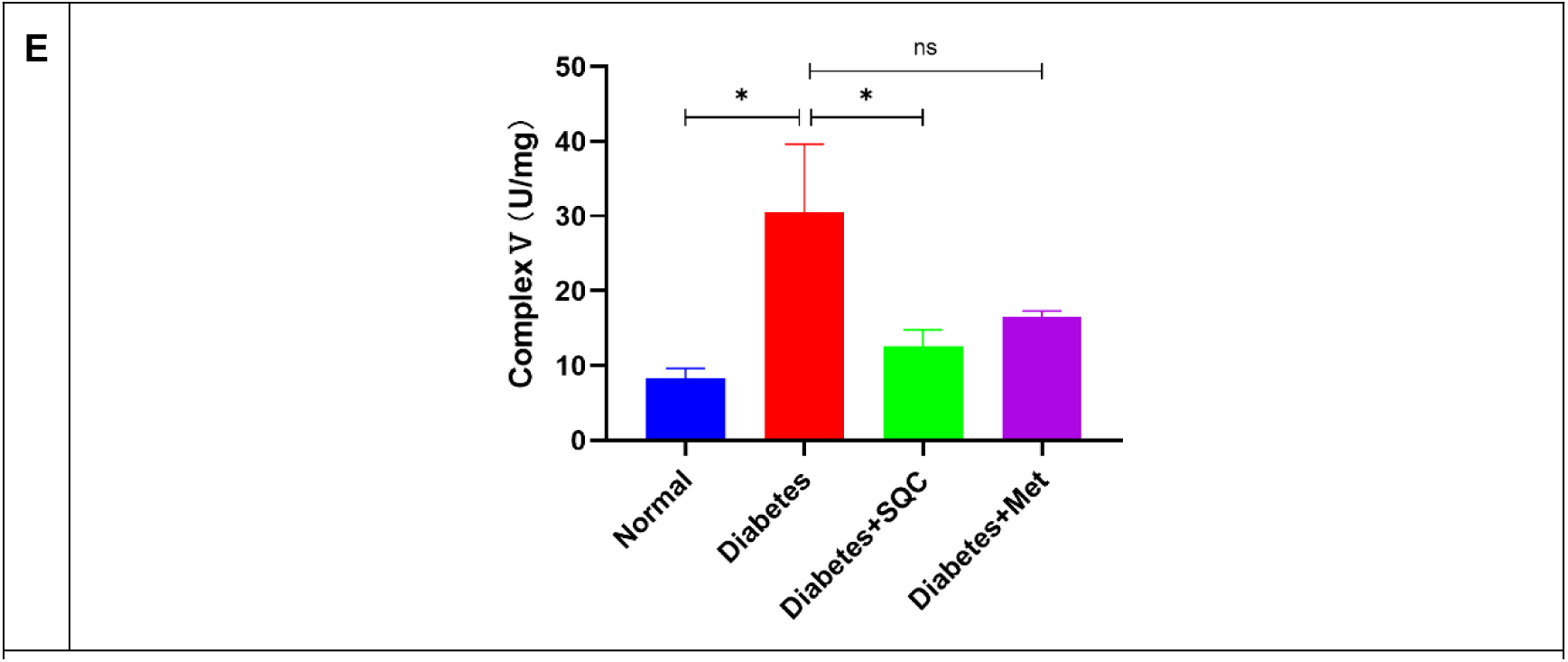
Effects of SQC and Met on mitochondrial complexes in pancreatic tissues of GK rats at 5-fold dilution. (A) Effect of SQC on complex I of GK rats, (B) Effect of SQC on complex II of GK rats, (C) Effect of SQC on complex III of GK rats, (D) Effect of SQC on complex IV of GK rats, (E) Effect of SQC on complex V of GK rats, and all the data were expressed as the mean ± SD (n = 3). * p < 0.05, ** p < 0.01.Abbreviation: NS, not significant.

### Effects of SQC on oxidative stress indices and ATP in rat pancreatic tissue

Pancreatic β-cell mitochondria are the main site of oxidative stress in cells, ROS, SOD, GSH are oxidative stress sensitive markers, if the mitochondrial function is damaged, these indexes will be changed, so ROS, SOD, GSH were measured, the results showed that ROS level in the model group was significantly higher than that of the control group (P<0.01), and the levels of SOD, GSH in the model group were significantly lower than that of the control group (P<0.05, P<0.01), however this relationship was reversed after SQC and Met treatment (Figure 5A-C). In addition, to further assess mitochondrial function, ATP, an indicator of mitochondrial energy metabolism, was measured, and it was found that ATP levels were significantly higher in the model group than in the control group (P<0.01), whereas ATP levels were significantly lower in the SQC and Met groups (Fig. 5D, P<0.05, P<0.01); thus, our results suggest that SQC and Met ameliorate the mitochondrial damage caused by high glucose.

**Figure 5.**
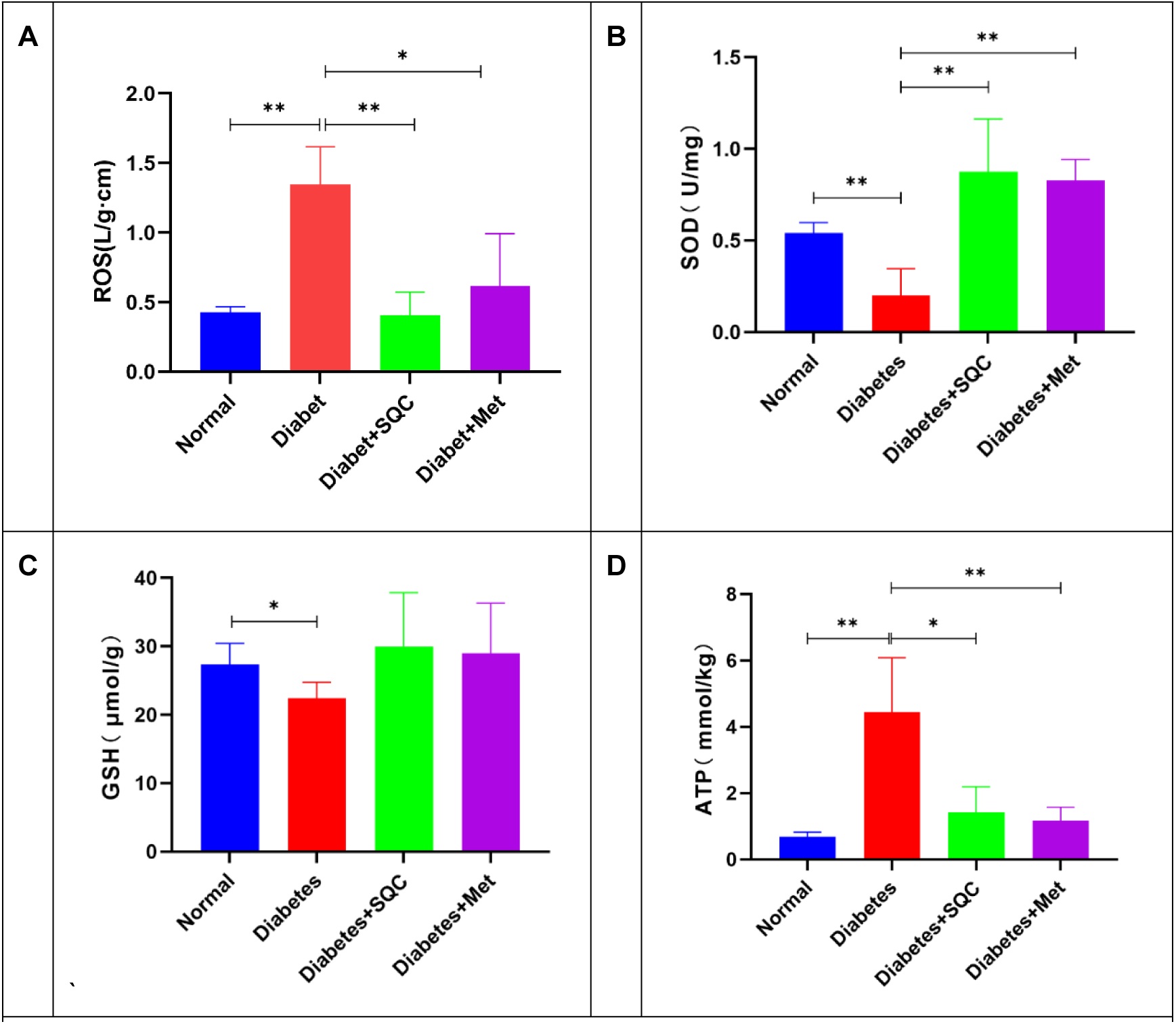
Effects of SQC and Met on ATP, ROS, GSH and SOD in pancreatic tissues of GK rats,.(A) Effect of SQC on ROS in GK rats, (B) Effect of SQC on SOD in GK rats, (C) Effect of SQC on cGSH in GK rats, (D) Effect of SQC on ATP in GK rats, All data are expressed as mean ± SD (n = 4). * P < 0.05, ** P < 0.01.

### Effect of SQC on large UCP-2 mRNA and protein

UCP-2, a transporter protein located on the inner mitochondrial membrane, is thought to be a major link between pancreatic β-cell dysfunction and T2DM. Therefore, we measured the expression levels of UCP-2 mRNA and protein using real-time PCR and protein blotting and analysis. Our results found that UCP-2 mRNA and protein expression was significantly elevated in the model group compared with the control group (Figure 6A-C,P<0.01), and UCP-2 mRNA and protein expression was significantly downregulated after 12 weeks of SQC and Met treatment (Figure 6A-C, P<0.05, P<0.01).

**Figure 6.**
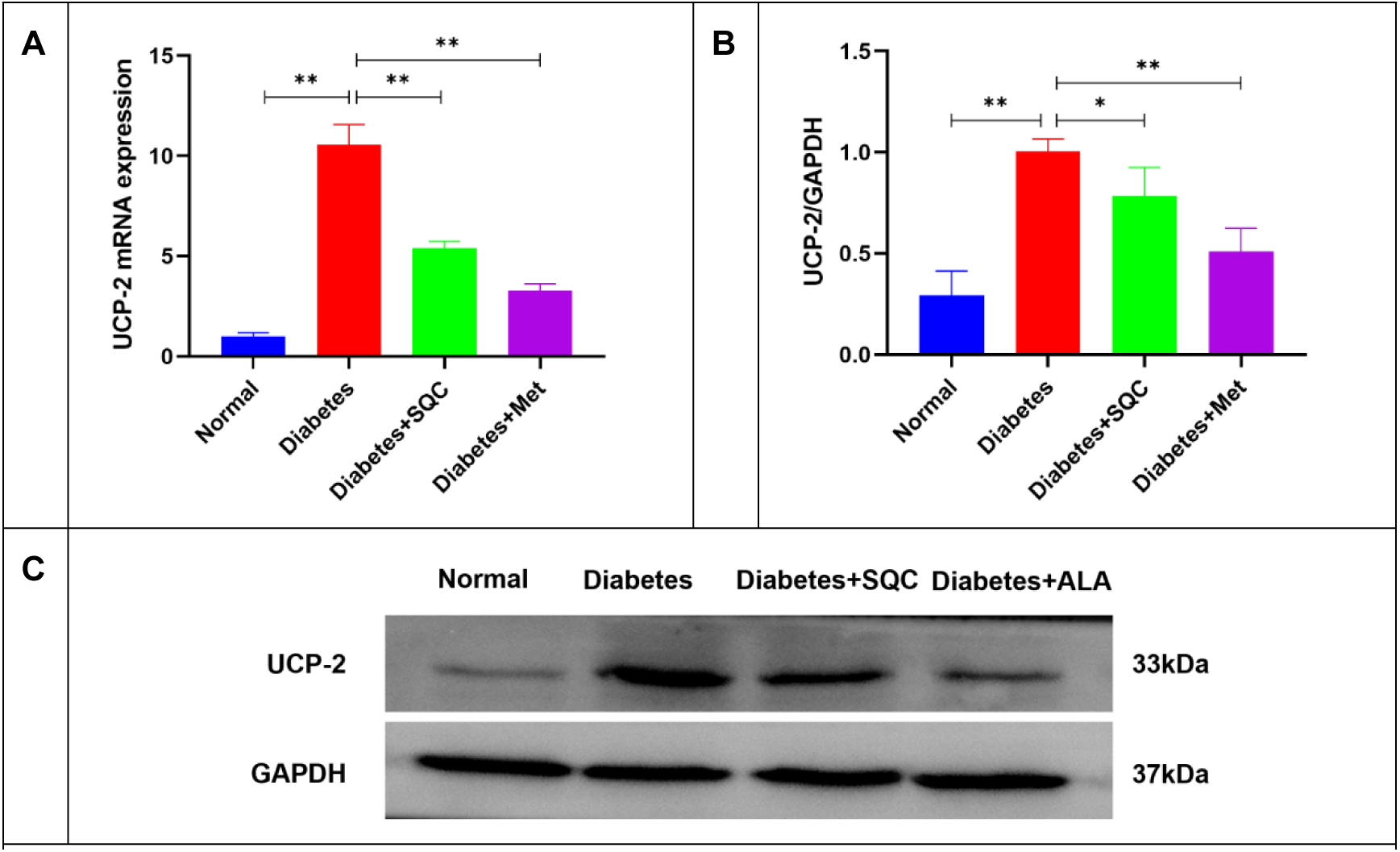
Effects of SQC and Met on UCP-2 in pancreatic tissues of GK rats, (A) Effects of SQC and Met on UCP-2 mRNA in GK rats, (B, C) Effects of SQC and Met on UCP-2 protein expression in GK rats, all data are expressed as mean ± SD (n = 4). * p < 0.05, ** p < 0.01.

## Discussion

T2DM is a metabolic disorder marked characterized by insulin resistance and defective pancreatic β-cell function. Insulin resistance is a typical manifestation of T2DM, but the progressive aggravation of insulin secretion malfunction due to the progressive decline of pancreatic β-cell function is the key factor in the progression of type 2 diabetes mellitus, and it is also an important target for diabetes mellitus prevention and treatment. The question of how to preserve pancreatic β-cells against harm while maintaining structural and functional integrity has become a clinical emphasis, although there is yet no successful applicable strategy.

In the process of preventing and treating T2DM, Chinese medicine has a long history of success. Its benefits include multi-targeting, significant efficacy, and few side effects. SQC is an empirical formula developed by Prof. Xie Chunguang, a national Qihuang scholar, and contains ginseng, astragalus, cornelian cherry meat, yam, danshen, raw rhubarb, smallpox pollen, and systemic rhubarb. It has been used in clinical settings for more than 20 years and clearly has islet-protecting properties. It has been clinically applied for more than twenty years and has a clear islet-protecting effect [30]. The benefits of SQC are most apparent in the protection of islet cells, where a large number of prior studies have shown that it reduces insulin resistance, inhibits the apoptosis of pancreatic islet-cells, controls glucose and lipid metabolism, and improves islet function in GK rats [19–21]. However, its specific role has not been fully elucidated, so the current study was carried out to further explore the molecular mechanism of this effect.

The progressive deterioration of insulin secretion due to progressive hypoplasia of pancreatic islet cell function is divided into four main stages: (1) compensatory stage: this stage is triggered by insulin resistance with compensatory hypertrophy of pancreatic islet β-cells, and insulin secretion is increased to counteract the insulin resistance in order to maintain glycemic stabilization; (2) mildly dysregulated stage: this stage exhibits fasting glucose abnormality due to the insufficiency of insulin secretion to compensate for the insulin resistance, glucose tolerance down-regulation, this stage of the pancreatic β-cell reserve function is reduced, insulin secretion is reduced; (3) severe loss of compensation: this stage of the pancreatic β- cells and structure seems normal, but its ability to synthesize insulin has been significantly reduced; (4) loss of compensation: this stage of the pancreatic islets appears obvious fat deposition/fibrosis, amyloidosis and other pathological structural changes, the pancreatic β- cells are significantly reduced in number and the loss of function [31]. As a result, of this experiment, the model group rats showed a significant increase in the level of FINS and a significant increase in FBG compared with the normal group rats, indicating that the pancreatic islets of the model groups were still in the stage of mild loss of compensation, and SQC could reduce the level of FINS in the diabetic GK rats, alleviate the pressure of insulin secretion in them, This also confirms the previous findings that SQC has the effect of improving insulin resistance and islet function; from the pathological results, SQC can improve the pathological damage of pancreatic tissue in diabetic GK rats, which also confirms the previous findings Modern research has demonstrated that glyco-lipotoxicity may trigger and aggravate progressive pancreatic-cell decompensation [32]; long-term elevated blood glucose, and “glyco-toxicity,” is the primary factor in pancreatic-cell failure [33], and “glyco-toxicity” results in progressive-cell decline due to decreased proliferation, increased apoptosis, and disrupted differentiation and dedifferentiation processes [34]; Lipotoxicity is another important triggering factor for type 2 diabetes, and when the level of free fatty acids in the blood exceeds the body’s carrying capacity, the FFAs are converted into triglycerides. Fatty acids (FFAs) are deposited as triglycerides (TG) in insulin-target tissues, resulting in “lipotoxicity.” FFAs, on the one hand, induce and exacerbate insulin resistance in peripheral tissues by promoting inflammatory responses; on the other hand, they act as a “prop” that directly destroys the structure and function of pancreatic islet cells, affecting insulin signaling pathways, as well as pancreatic islet cell proliferation and apoptosis [35]. However, glucotoxicity and lipotoxicity do not act independently to impair islet cell function; rather, they work in a synergistic manner. Together, glycotoxicity and lipotoxicity, often known as glycolipotoxicity, facilitate the gradual loss of islet cell structure and function [36]. In the present research, it was discovered that both SQC and Met could efficiently lower the FBG level in GK rats through biochemical testing of blood glucose and lipids; however, SQC’s effect on lowering blood glucose was smoother. In addition, it was discovered that SQC and Met could significantly lower the TC, TG, and LDL levels as well as elevate the HDL-C levels of diabetic rats;This also confirms previous findings that SQC has improved glycolipid metabolism.

The primary location of energy metabolism in Pancreatic β-cell is the mitochondria, and the damaging impacts of glucolipotoxicity on pancreatic islet cells are first recognized by increasing metabolic stress on the cells. Under high-glucose stimulation, glucose stimulation in mitochondria is primarily accomplished through the Tricarboxylic Acid Cycle pathway (TCA) pathway for ATP formation, and the other is the electron transport chain pathway, which is the pathway of nicotinamide reduction by reduced nicotinyl acetate (NADH). The other is the electron transport chain pathway, in which O_2_ is reduced by reduced nicotinoyl purine dinucleotide (NADH), ultimately resulting in the creation of H_2_O. The electron transfer order is complex I coenzyme Q complex III complex IV. The electron donor in the second electron transport chain pathway is succinate FADH II, and the electrons are transferred in the following order: complex II, coenzyme Q, complex III, and complex IV. Numerous protons (H^+^) are present beside the inner mitochondrial membrane during this electron conversion process.

During this electron conversion process, the inner mitochondrial membrane is accompanied by the generation of a large number of protons (H^+)^, in which protons are pumped into the mitochondrial membrane gap from the mitochondrial matrix by the mitochondrial membrane with the proton pumping function of the complex (COMPLEXI, III, IV). The free radicals released at this stage can induce adenosine diphosphate (ADP) to generate a large amount of ATP by coupling with Pi under the influence of complex V, which on the one hand provides energy for the cells, and on the other hand acts as a signaling factor to regulate insulin secretion [37,38]. The results of the current study indicated that SQC down-regulated the expression of mitochondrial respiratory chain complex I-V and reduced its ATP formation, suggesting that SQC could alleviate the metabolic stress in pancreatic tissues.

UCP-2 is the main intracellular ROS reduction mechanism and is believed to have an essential function in the adverse regulation of insulin secretion [39]. Normally, metabolized ROS are swiftly eliminated up by the cellular antioxidants SOD and GSH [40], repairing damaged cells and reducing organism damage. However, under the stress of energy metabolism caused by long-term glycolipotoxicity, excess glucose or fatty acids produce accelerated ROS production, leading to ROS accumulation [41,42]. ROS enable insulin and glucagon to be damaged by triggering intracellular inflammatory responses, hence mediating islet dysfunction. ROS disrupt cell structure and function by activating intracellular inflammatory responses, mediating pancreatic-cell apoptosis, and directly destroying cellular proteins, among other things. UCP- 2 acts as a critical link in the regulation of ROS and, as a result, pancreatic islet function. By returning hydrogen ions from the respiratory chain to the mitochondrial matrix via the inner mitochondrial membrane, it can neutralize ROS [39]. UCP-2 can, however, interfere with ATP generation when it is overexpressed by altering the membrane potential in the electron transport chain [45]. The results of the present research indicated that SQC downregulated the gene and protein expression of UCP-2 mRNA, which could effectively prevent the overexpression of UCP-2, increased the expression of antioxidants SOD and GSH, reduced the accumulation of ROS, and raised the expression of SOD and GSH. The effects of UCP-2 on ATP synthesis and insulin secretion in rat pancreatic tissues were not evident since the GK animals in each model group in our experiment were still in the compensatory stage.

There are some limitations of this study, firstly, this study only investigated the protective effect of SQC on the function of islet cells in the compensated stage due to laboratory limitations, the protective effect of SQC on the function of islet cells in the decompensated stage has not yet been investigated; previous reports have found that mitochondrial dysfunction is an important mechanism of T2DM-induced muscle damage, and that elevated free fatty acids in the plasma of patients with T2DM and inflammatory cytokines cause impaired oxidative capacity of muscle mitochondria, leading to decreased fat oxidizing activity and increased intracellular lipids in muscle cells, which in turn causes abnormal energy metabolism in muscle cells [46]. In this study, only the results of SQC on mitochondrial energy metabolism in pancreatic tissues were investigated, and the effects on skeletal muscle were not further observed; moreover, in terms of mechanism study, only a single index was selected in this experiment to investigate the effects of SQC on protein and gene expression of UCP-2 in pancreatic tissues, Next, proteomic and transcriptomic studies should be conducted on the target genes and proteins of key pathways in pancreatic and skeletal muscle tissues to fully evaluate the mechanism of protective effects of SQC on mitochondrial energy metabolism.

## Conclusion

In conclusion, SQC can improve glucose-lipid metabolism levels and pancreatic histopathology in diabetic GK rats, exerting a protective effect on pancreatic islet cells. Its main mechanism of action may be to relieve the metabolic stress of pancreatic islet mitochondria by inhibiting oxidative stress

## Acknowledgments

We would like to acknowledge the hard and dedicatedwork of all the staff that implemented the interventionand evaluation components of the study.

## Data availability statement

Data associated with our study has not been deposited into a publicly available repository. All data generated or analyzed during this study are included in this article and its supplementary information files.

## Ethics statement

This study was performed according to the guidelines and recommendations of the Experimental Animal Ethics Committee of the Affiliated Hospital of Chengdu University of Traditional Chinese Medicine, Sichuan Province, China, review number: 2021DL-011.

## statement

I promise that all data will be open and accessible to everyone.

